# Balancing Barcoding and Genomics: gDNA Quality in Insect Vouchers after HotSHOT DNA extraction

**DOI:** 10.1101/2025.10.09.680689

**Authors:** Vivian Feng, Caroline Høegh-Guldberg, Aleš Buček, Rudolf Meier

## Abstract

DNA extraction with an alkaline buffer system called “HotSHOT” is widely used for barcoding because it is rapid, inexpensive, and voucher preserving, but it remains unclear whether sufficient genomic DNA (gDNA) remains in small vouchers for downstream use in genomics. We here evaluate gDNA quality and quantity before and after HotSHOT treatment of 11 insect families representing six orders. Some specimens were flash frozen immediately after collection, while others were kept for one week at tropical temperatures in ethanol to mimic Malaise trap conditions. Encouragingly, we show that gDNA of sufficiently high quality and quantity for genomic sequencing remained in specimens treated with HotSHOT. We also show that DNA integrity was strongly influenced by field storage with specimens exposed to Malaise trap conditions showing such pronounced degradation that the standard HotSHOT treatment no longer significantly altered DNA quality. For control material, HotSHOT treatments involving longer exposure to high temperature led to smaller fragment lengths with the effect apparently being influenced by the degree of specimen sclerotization. Our results thus suggest that optimized HotSHOT treatments, together with carefully controlled pre-extraction storage, preserve voucher gDNA of sufficient quality for downstream genomic analyses with both short-read and possibly even some long-read sequencing technologies. Our protocol selection guidelines improve voucher gDNA preservation in HotSHOT-treated samples. This is particularly important for many species which are only known from one or few specimens discovered during barcoding projects.

## Introduction

Bulk sampling methods such as Malaise, pitfall, and light traps routinely collect thousands of specimens belonging to hundreds of species in a single event (Srivathsan et al., 2023). Such samples can be thought of as potential metagenomic libraries in which genetic diversity is largely partitioned by species because intraspecific variability is generally much smaller than interspecific variability. Because many species are singletons and only a few dominate in numbers (see fig 1 in Meier et al., 2024), performing full DNA extractions and sequencing for every individual is neither efficient nor desirable for answering most biological questions. It concentrates effort and budget on repeatedly sequencing common species, and it creates a “two-voucher problem” in that every specimen generates not only a physical voucher but also a separate DNA extract vial that must be archived, tracked, and curated. This duplicates storage needs, inflates consumables and freezer costs, complicates chain-of-custody, and can compromise morphological integrity of specimens when aggressive DNA extraction techniques are used. Many scientists thus use a different workflow that starts with generating specimen-specific barcodes that are then used for rapid sorting to approximately species level. In the next step, only selected specimens are subjected for targeted downstream analyses to answer specific questions. Such a “reverse workflow” (Wang et al., 2018), where barcoding precedes morphological or genomic exploration of select specimens, has been demonstrated to be technically and financially feasible due to the wide availability of third-generation sequencing platforms (Hebert et al., 2018; Srivathsan et al., 2018, 2019, 2021, 2024). Such workflows are thus also considered central to accelerating biodiversity discovery at scale to address ‘entomological dark matter’ (Hartop et al., 2024; Chua et al., 2023; Brydegaard et al., 2024; Meier et al., 2024, 2025a).

**Figure 1.**
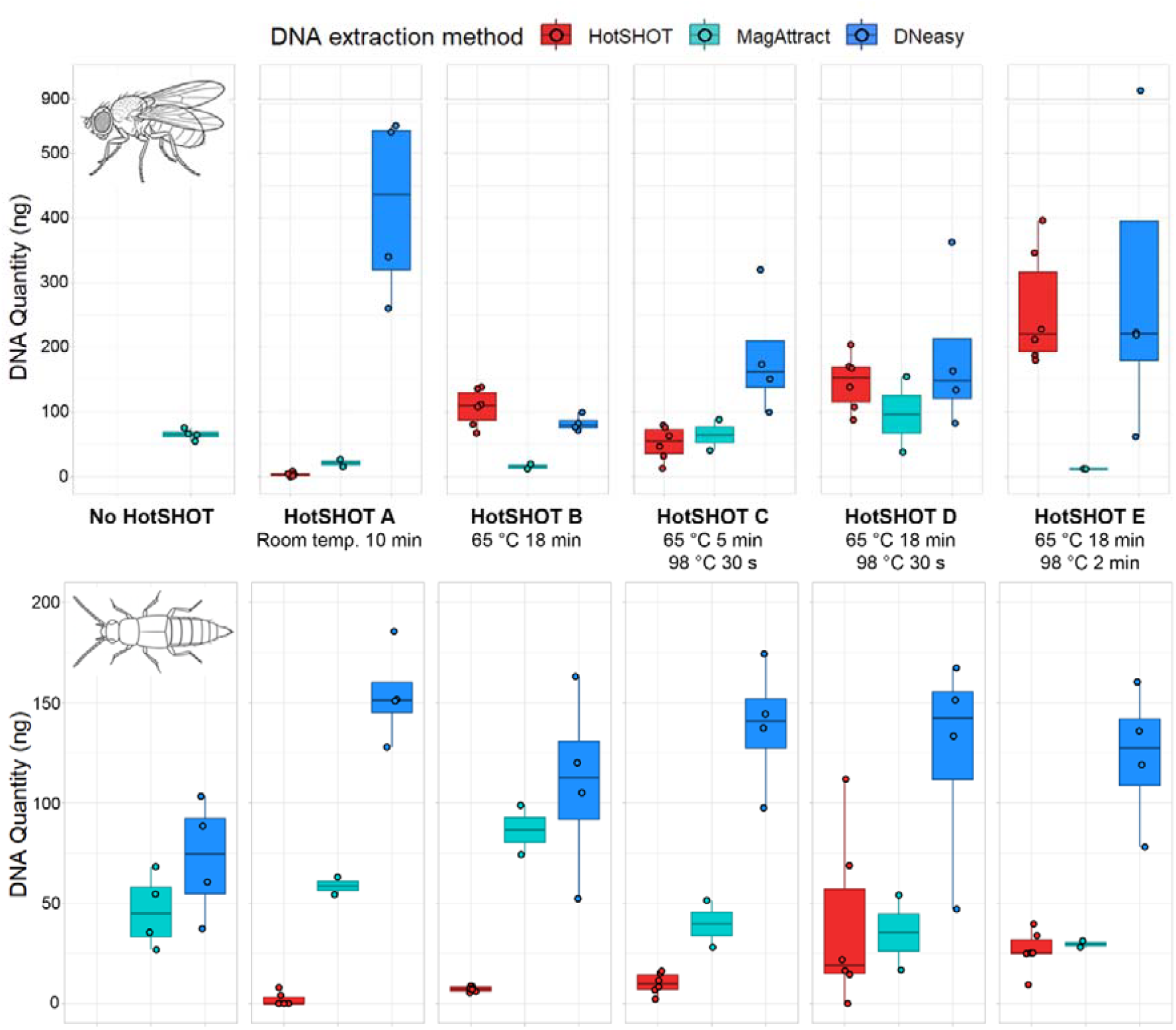
Increasing intensity of HotSHOT treatment increases DNA quantity in HotSHOT extract but does not affect DNA quantity in subsequent genomic extractions (with MagAttract and DNeasy kits) of the HotSHOT-treated specimens.

Data produced during this two-step workflow can directly or indirectly address all seven biodiversity shortfalls highlighted in Hortal et al. (2015). The Linnean shortfall of undescribed species is reduced because barcodes can be used for species discovery in bulk samples and preparing them for integrative taxonomy (Hartop et al., 2022; Amorim et al., 2025; Meier et al., 2025a). The Wallacean shortfall, concerning incomplete distribution knowledge, and the Prestonian shortfall, concerning missing abundance data, are also addressed, because each barcoded specimen is counted and linked to a collection locality. Unfortunately, the remaining biodiversity shortfalls in Hortal et al. (2015) require more expensive, genomic data (Luikart et al., 2003). The Darwinian shortfall, incomplete phylogenies, can be tackled by obtaining low-coverage genomes for barcoded vouchers as long as they yield enough phylogenetically informative markers (e.g., Call et al., 2023). The Raunkiæran shortfall on traits can be reduced by combining morphological information with genomic data on functions such as metabolic pathways. The Hutchinsonian shortfall on abiotic tolerances can be approached by examining population genomic variation for signals of adaptation, while the Eltonian shortfall on ecological interactions can occasionally already be addressed with the DNA obtained in the first step (Quintana et al. 2022; Srivathsan et al., 2022) but usually requires community-level genomic datasets.

A critical element for the feasibility of this two-step pipeline is that vouchers retain genomic DNA (gDNA) of sufficient quality and quantity after barcoding. This explains the popularity of techniques that just “leaches” a small amount of DNA from the specimen. Several methods have been proposed (e.g., Margam et al., 2010; Wong et al., 2014; Thongjued et al., 2019; Wang et al., 2018), but many labs have recently started to use HotSHOT because it is rapid, inexpensive, and preserves vouchers (Truett et al., 2000). HotSHOT extraction skips DNA purification and only requires two inexpensive buffers, one for leaching DNA and one for neutralization (Srivathsan et al., 2023; Fikáček et al., 2024). Neither needs to be kept cold. Unfortunately, however, it remains unclear how much DNA remains in the specimens after HotSHOT extraction. Work on microbiomes after HotSHOT leaching has provided mixed signals (Andriienko et al., 2024): the overall DNA yield was reduced and the fragment length distribution shifted towards smaller DNA fragments. Yet, it did not appear to bias the reconstructed bacterial communities when assessed with 16S sequencing. This suggests that HotSHOT may not alter DNA quality and quantity to such an extent that it would prevent meaningful downstream genomic analyses based on short amplicon sequencing.

Here, we explicitly tested whether insect vouchers treated with HotSHOT retain gDNA of sufficient integrity for low-coverage genome sequencing. We carried out two experiments. We quantified gDNA of fresh control specimens for two species whose DNA was extracted with and without exposure to HotSHOT. In the second experiment, we asked whether DNA degradation caused by HotSHOT is more limiting than the degradation experienced while specimens are in a Malaise trap for one week in the tropics. To test this, we exposed a taxonomically diverse set of specimens to conditions typical of Malaise traps pitched in a tropical country. We then compared gDNA quality and quantity of specimens with and without exposure to HotSHOT. Our results serve as empirical guidelines for firstly assessing and then maximizing the quantity and quality of gDNA remaining in the specimen post-extraction.

## Materials and Methods

### Control: Impact of HotSHOT on DNA quality and quantity in control specimens

Fresh *Drosophila melanogaster* (Diptera: Drosophilidae) from a laboratory colony and freshly collected *Myllaena dubia* (Coleoptera: Staphylinidae) from the Slavkov Forest, Czech Republic (50.054N, 12.859E) were flash frozen in liquid nitrogen and stored in absolute ethanol at −80 °C prior. The gDNA of control specimens was extracted immediately without HotSHOT (Truett et al., 2000) exposure, thus providing the baseline for expected gDNA quality and quantity. The remaining specimens were exposed to different HotSHOT treatments. The first was leaving specimens in HotSHOT at room temperature for 10 min before extraction (“A”). The next treatment applied the standard HotSHOT treatment (65 °C for 18 min plus 98 °C for 2 min; “E”), which is widely used in high throughput barcoding workflows, but we also tested three additional HotSHOT treatments with shorter exposure: treatment “B”, 65 °C for 18 min; treatment “C”, 65 °C for 5 min plus 98 °C for 30 s; and treatment “D”, 65 °C for 18 min plus 98 °C for 30 s. To evaluate the effect of HotSHOT exposure, the gDNA of all specimens was extracted with either the Qiagen DNeasy Blood and Tissue Kit (non-destructive extraction, final elution into 25 µl of water; n = 4 biological replicates) or Qiagen MagAttract HMW DNA Kit (liquid nitrogen–frozen individuals disrupted with mortar and pestle, final elution into 20 µl of water; n = 2–4 biological replicates). The MagAttract Kit-based DNA extraction followed modified manufacturer’s protocol with following modifications: the specimen frozen by liquid nitrogen was pulverized with plastic pestle in 1.5 ml plastic tube and 187 µl of mastermix (consisting of 100 µl PBS buffer, 10 µl of Proteinase K, 2 µl of RNase A, and 75 µl of Buffer AL) was added to the still frozen pulverized sample and was incubated at room temperature for 2 hours with gentle inversion of the tube every 30 minutes. All subsequent incubation steps except for the last elution step were performed without shaking at room temperature. The Blood and Tissue Kit-based DNA extraction followed modified manufacturer’s protocol with following modifications: the specimen was submerged in 180 µl of ATL buffer and 20 µl Proteinase K mix, vortexed briefly for 15 seconds and incubated overnight at 56 °C without shaking. This protocol did include physical sample disintegration and homogenization, and the extracted specimens are typically intact with occasional breakage of some appendages during the procedure.

### Malaise Trap: Impact of HotSHOT on DNA quality and quantity of exposed specimens

We next asked whether typical delays between a specimen falling into a Malaise trap collecting jar and its gDNA being extracted have a strong impact on DNA quality before and after HotSHOT treatment. To test this, specimens from 9 families representing 5 insect orders and spanning a wide range of body sizes were collected at Kent Ridge Park, Singapore (1.287N, 103.789E) and stored in a Malaise trap jar filled with 70% ethanol (Diptera: Sepsidae; Hemiptera: Alydidae, Cicadellidae, Pyrrhocoridae; Hymenoptera: Apidae; Lepidoptera: Lycaenidae; Orthoptera: Gryllidae). The jar was left outdoors in a partially shaded environment for one week to mimic tropical field conditions. After this period, half the specimens for each family were extracted directly with the Qiagen DNeasy Blood and Tissue Kit (final elution into 50 µl of buffer AE), while the other half were first subjected to the standard HotSHOT treatment (“E”: 65 °C for 18 min, 98 °C for 2 min; buffer volume 15–20 µl) and then extracted with the same kit. Entire insects were processed for most families, while only mid legs were used for the larger specimens of Apidae and Gryllidae.

### DNA quality assessment with Qubit and Tapestation

The DNA concentration of each extract (from HotSHOT, DNeasy and MagAttract) was measured with the Qubit fluorometer. Purified gDNA extracts (from DNeasy and MagAttract) were assessed with the Genomic DNA ScreenTape (Agilent) assay for DNA integrity, given by the DNA Integrity Number (DIN). Fragment length distributions were obtained by classifying the most common fragment sizes of each sample, given in tables under each electropherogram, into one of five size classes (more than 10 kbp, 5 – 10 kbp, 1 – 5 kbp, 300 bp – 1 kbp, less than 300 bp). Estimated quantities of each fragment size group were calculated by multiplying integrated areas provided in the same table with sample DNA concentrations from Qubit. Both DINs and electropherograms were obtained via the TapeStation Analysis Software (versions A.02.01 and 5.1).

### Amplification

Because varying intensity and duration of HotSHOT treatments can have a measurable impact on the quality and concentration of the HotSHOT extract (hsDNA), we also tested for the fresh specimens whether the hsDNA still contained sufficient template DNA to yield reliable PCR products. We attempted to amplify the Folmer region (658 bp) of the COI gene from the hsDNA using indexed LCO1490 and HCO2198 primers (Folmer et al., 1994). The primers had 9 bp indices attached at the 5’ end, sourced from Srivathsan et al. (2024), allowing for subsequent demultiplexing. PCR reactions consisted of 10 µl hsDNA, 9 µl PCR water, 1 µl forward and reverse primer, and 4 µl HOT FIREPol Blend Master Mix (04-25-00120-10, Solis BioDyne). Cycling conditions were: 94 °C for 15 min, followed by five cycles (94 °C for 30 s, 47 °C for 40 s, 72 °C for 1 min), 30 cycles (94 °C for 30 s, 52 °C for 40 s, 72 °C for 1 min), and a final extension of 72 °C for 7 min. Amplicons were visualized by electrophoresis on a 2% agarose gel.

### Statistical analyses

All analyses were performed in R version 4.4.2 (R Core Team, 2024) using RStudio version 2023.06.0 (Posit team, 2023). DNA yield and integrity were analyzed with analysis of variance (ANOVA) models fitted using the function *aov()*. Post-hoc multiple comparisons among HotSHOT treatments were carried out with Tukey’s HSD test using the function *TukeyHSD()*. To summarize groupings, compact letter displays were generated with the function *multcompLetters4()* from the *multcompView* package (Graves et al., 2019), where groups sharing a letter are not significantly different. For the Malaise trap sample, DIN values were compared using Welch’s two-sample *t*-test with the function *t*.*test()*. Figures were produced with the *ggplot2* package (Wickham, 2016). Boxplots were drawn with *geom_boxplot()*, where the central line indicates the median, the box spans the interquartile range (IQR) and whiskers extend no further than 1.5 × IQR. Individual observations were overlaid as jittered points with *geom_jitter()*. For line plots, group means were added with *stat_summary(fun = mean, geom = “point”)* and connected with *stat_summary(fun = mean, geom = “line”)*. Error bars were calculated as ± standard error of the mean using stat_summary(fun.data = mean_se, geom = “errorbar”).

## Results

### Impact of HotSHOT on DNA quality and quantity in control specimens

The amount of temperature exposure during HotSHOT influenced the amount of DNA in HotSHOT extract (hsDNA) of control specimens. The quantity of hsDNA was strongly dependent on the temperature protocol of HotSHOT. hsDNA yield increased with the temperature exposure during HotSHOT, with the standard treatment “E” producing the largest quantities (Fig. 1) as tested in a two-way ANOVA of a linear model modelling taxon-specific differences (Amount of hsDNA ∼ Treatment * Taxon). It revealed significant effects of treatment (F_4,50_ = 26.27, p < 0.001), taxon (F_1,50_ = 102.93, p < 0.001) and the interaction between treatment and taxon (F_4,50_ = 16.93, p < 0.001). Surprisingly, the amount of genomic DNA (gDNA) extractable from the HotSHOT-treated specimen for further genomic work was not significantly reduced when increasing high-temperature exposure during HotSHOT treatments (Fig. 1) as tested in a two-way ANOVA (Amount of gDNA ∼ Treatment * Taxon; F_5,60_ = 1.90, p = 0.108) nor the interaction (F_5,60_ = 1.09, p = 0.376) and the only significant effect was the variable “Taxon” with the amount of gDNA in the specimens being significantly higher in *D. melanogaster* than *M. dubia* (F_1,60_ = 4.11, p = 0.047). The DNeasy extraction method also produced significantly higher amounts of gDNA than the MagAttract extraction method (Amount of gDNA ∼ Treatment * DNA extraction method; F_1,68_ = 18.17, p < 0.001) but neither treatment (F_1,68_ = 0.91, p = 0.344) nor the interaction (F_1,68_ = 2.24, p = 0.139) was significant.

Across all treatments, *Drosophila melanogaster* consistently produced higher yields than the more heavily sclerotized *Myllaena dubia*. However, yield alone is not a reliable indicator of usable DNA, because treatments that maximize yield may also produce the most degraded extracts. This effect was confirmed in our experiments. Untreated control specimens had the highest gDNA quality as quantified by DIN, while even a brief 10-minute room temperature exposure to HotSHOT caused a measurable decline in quality (Fig. 2). DIN decreased further under all HotSHOT incubations, with the most widely adopted treatment “E” producing the lowest DIN. These results were statistically significant with a one-way ANOVA showing that treatment had a significant effect on DIN (DIN ∼ Treatment; F_5, 66_ = 15.69, p < 0.001) and a post-hoc Tukey HSD test showing significantly higher DIN for the control (group “a”) compared to all other treatments (groups “b” and “c”) except treatment “B” (group “ab”) and significantly lower DIN for treatment “E” (group “c”) compared to all other treatments (Fig. 2). Less sclerotized *D. melanogaster* had significantly higher DINs than more sclerotized *M. dubia* across treatments (DIN ∼ Treatment * Taxon; F_1, 60_ = 5.59, p = 0.021) except for treatment “E” where they exhibited a large decline in DIN (Fig. 4) which suggests greater resilience to high temperature treatments for more sclerotized taxa. In the same two-way ANOVA, both treatment (F_5, 60_ = 19.17, p < 0.001) as well as the interaction between treatment and taxon (F_5, 60_ = 3.01, p = 0.017) were significant.

**Figure 2.**
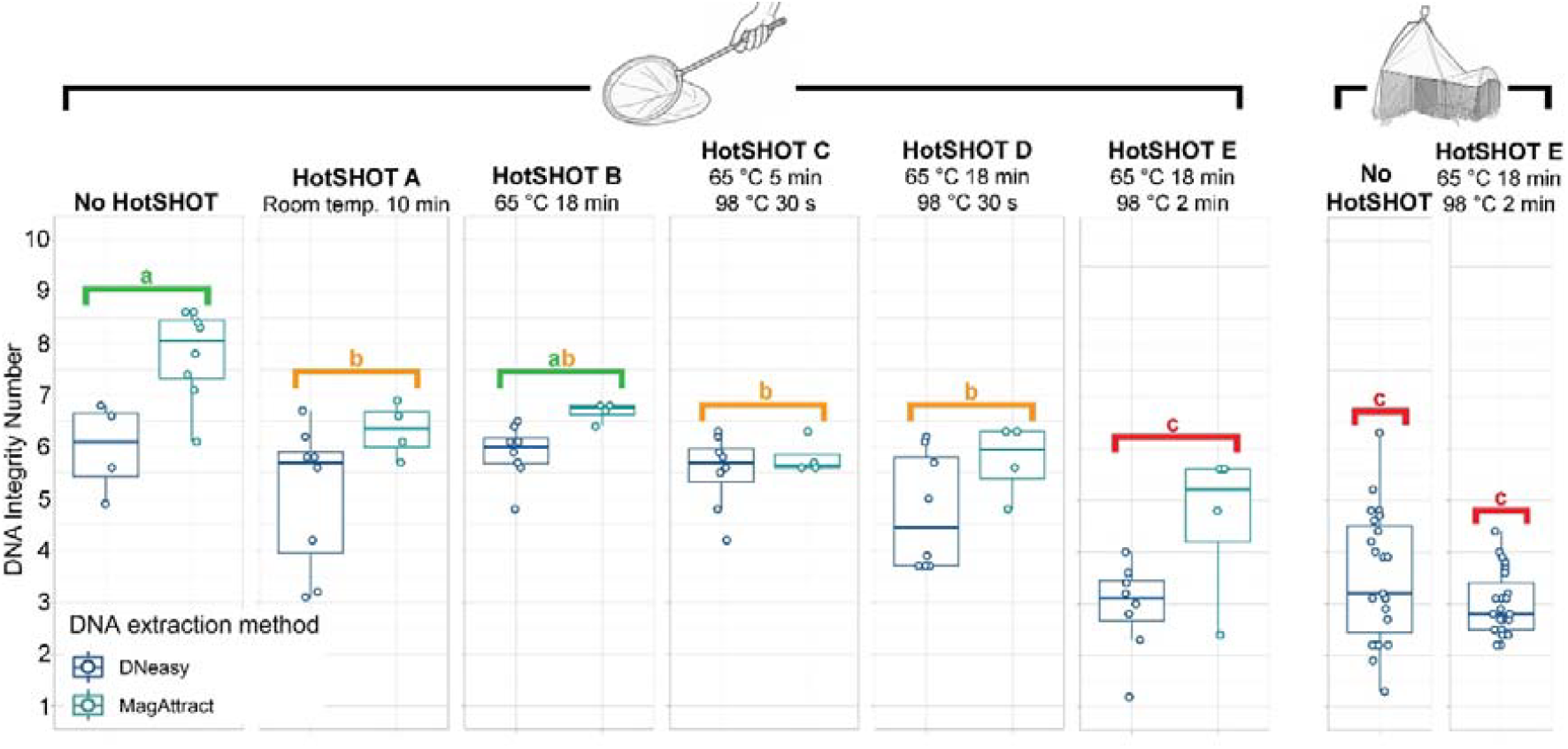
HotSHOT treatment caused a measurable decline in DNA Integrity Number (DIN) in control samples, with untreated specimens having significantly higher DIN than all treatments except “B”. HotSHOT treatment did not cause a significant decline in DIN in the Malaise trap sample. Groups sharing a letter above the boxplot are not significantly different from each other.

Size distributions of gDNA fragments provided a complementary perspective (Fig. 3). Untreated control specimens contained substantial amounts of high molecular weight gDNA, with fragments longer than 10 kbp being abundant. Specimens with HotSHOT treatments “B”, “C” and “D” showed a similar profile of gDNA fragments, though with greater variability in amount of fragments longer than 10 kbp. Under the standard HotSHOT treatment “E”, gDNA fragments larger than 10 kbp mostly disappeared and most gDNA fragments varied in length between 1 and 5 kbp. Note, however, that despite strong differences in yield and quality, PCR amplification of the COI barcode was robust across most HotSHOT treatments (Fig. 5) with amplification being successful for both taxa under all extraction treatments. However, band intensity was reduced for *M. dubia* in treatments lacking a 98 °C step.

**Figure 3.**
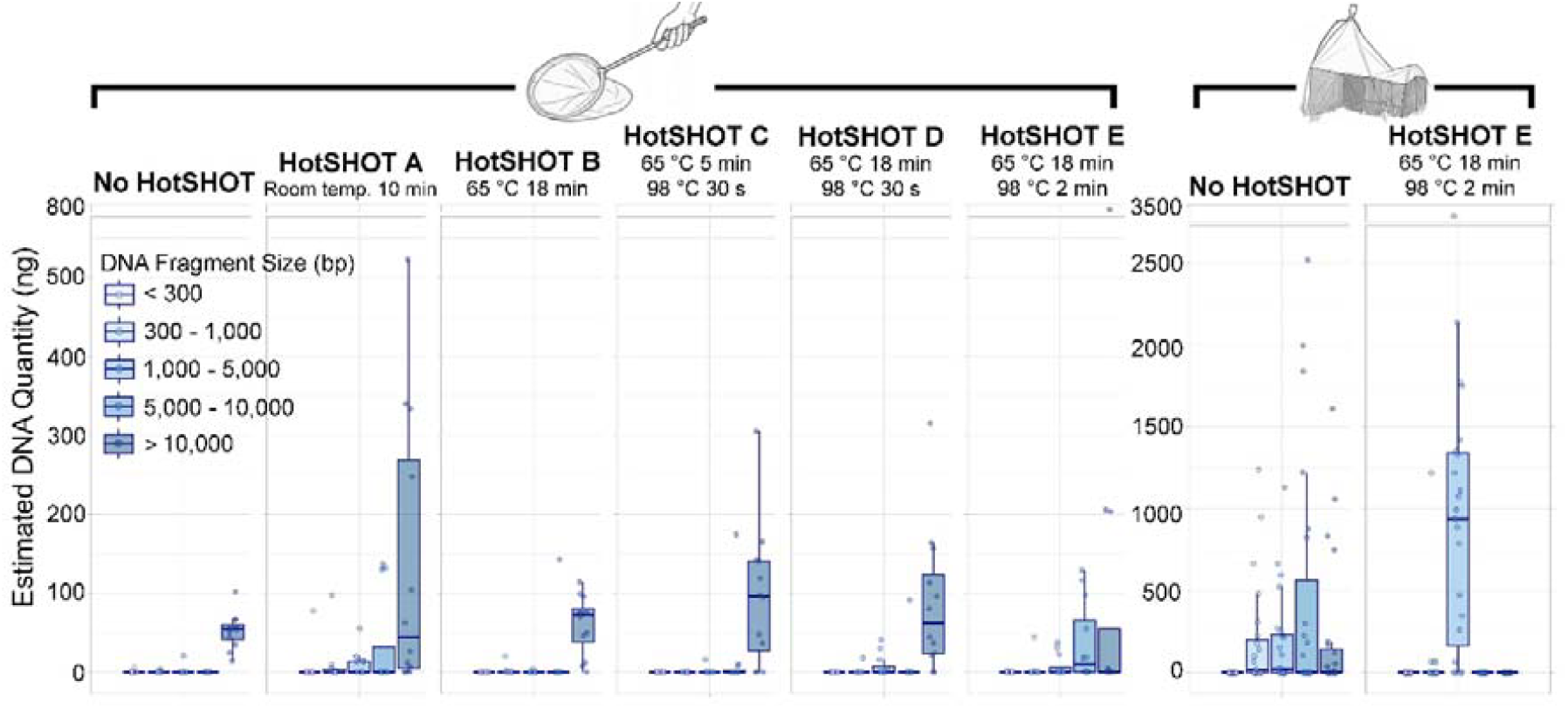
Only HotSHOT treatment “E” reduced the amount of fragments over 10 kbp in gDNA of control specimens obtained with both extraction kits. Using a Malaise trap results in more fragmented gDNA, and the combination of a Malaise trap and HotSHOT treatment “E” eliminated all remaining long fragments.

### Impact of HotSHOT on DNA quality and quantity of Malaise trap specimens

Overall, the Malaise trap experiment showed that extended exposure of specimens to field conditions was a major driver of DNA degradation, effectively being as destructive as the harshest HotSHOT treatment (roughly equivalent to treatment “E” in experiment 1; Fig. 2) thus leaving less scope for further decline after applying HotSHOT treatment “E” because most gDNA was already fragmented to 300 bp - 5 kbp size range and fragments above 10 kbp were rare (Fig. 3). Additional HotSHOT treatment shifted these distributions further, with treatment “E” eliminating nearly all remaining long fragments and making the 1–5 kbp size class dominant (Fig. 3). Accordingly, the mean DIN value for Malaise trap specimens after HotSHOT treatment declined only moderately from 3.26 to 3.00, a difference that was not significant (Welch’s two-sample t-test, 32.31 = 1.76, p = 0.088), while experiment 1 yielded a significant decline between control specimens (DIN=7.18) and specimens with HotSHOT treatment E (DIN=3.49). When we used a linear model for the taxa in the Malaise trap experiment, (DIN ∼ Treatment * Taxon), we found in a two-way ANOVA that both treatment (F_1, 30_ = 7.13, p = 0.012) and taxon (F_7,30_ = 8.91, p < 0.001) had a significant effect on DIN, but the interaction between treatment and taxon was not significant (F_7,30_ = 1.25, p = 0.309). This implied that the treatment effects were consistent across taxa (Fig. 4).

**Figure 4.**
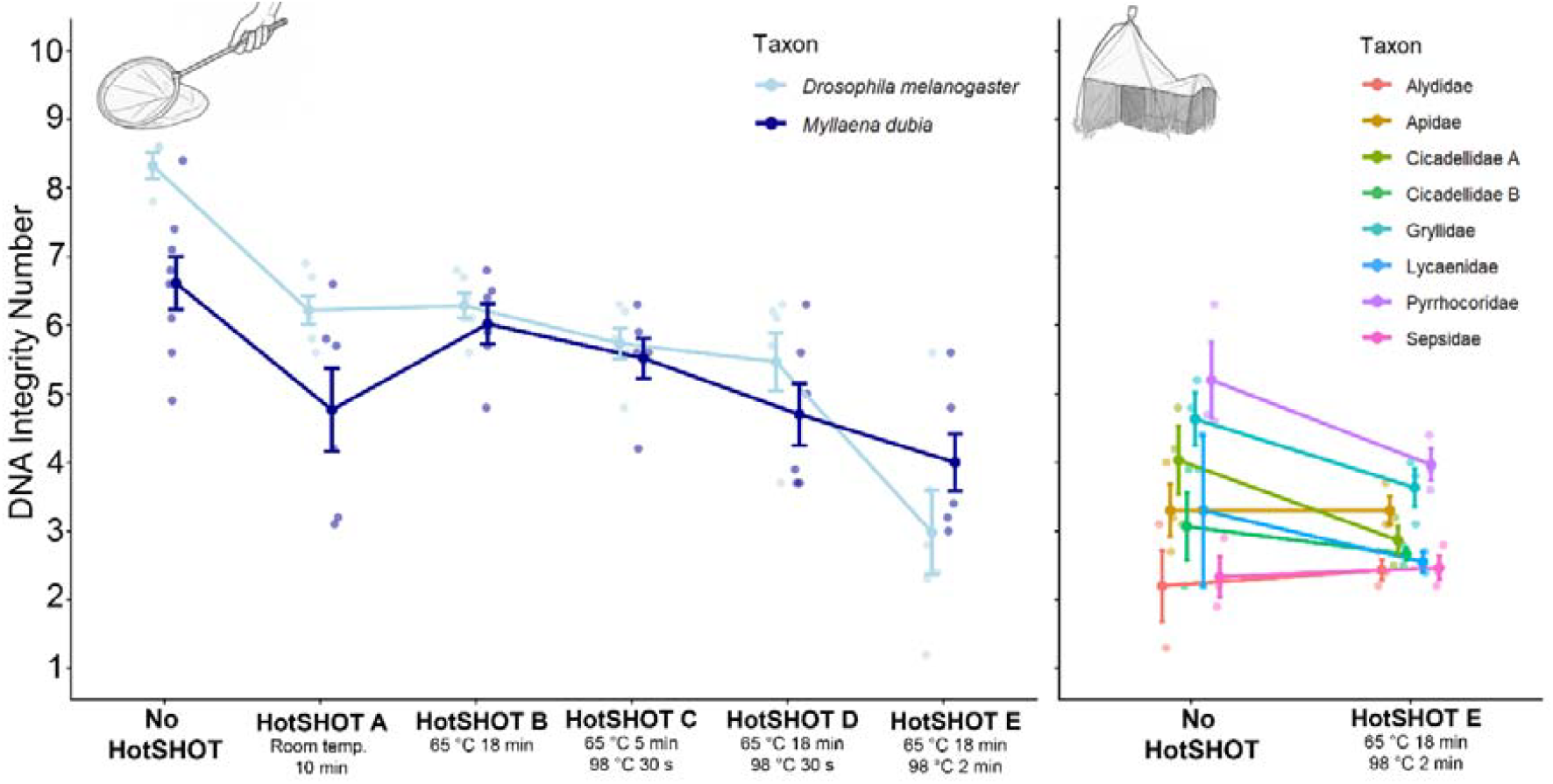
Less sclerotized taxa (e.g. *D. melanogaster*) have a stronger response than more sclerotized taxa (e.g. *M. dubia*) to the most intense HotSHOT treatment (“E”), while taxon did not have a significant effect in the Malaise trap sample.

**Figure 5.**
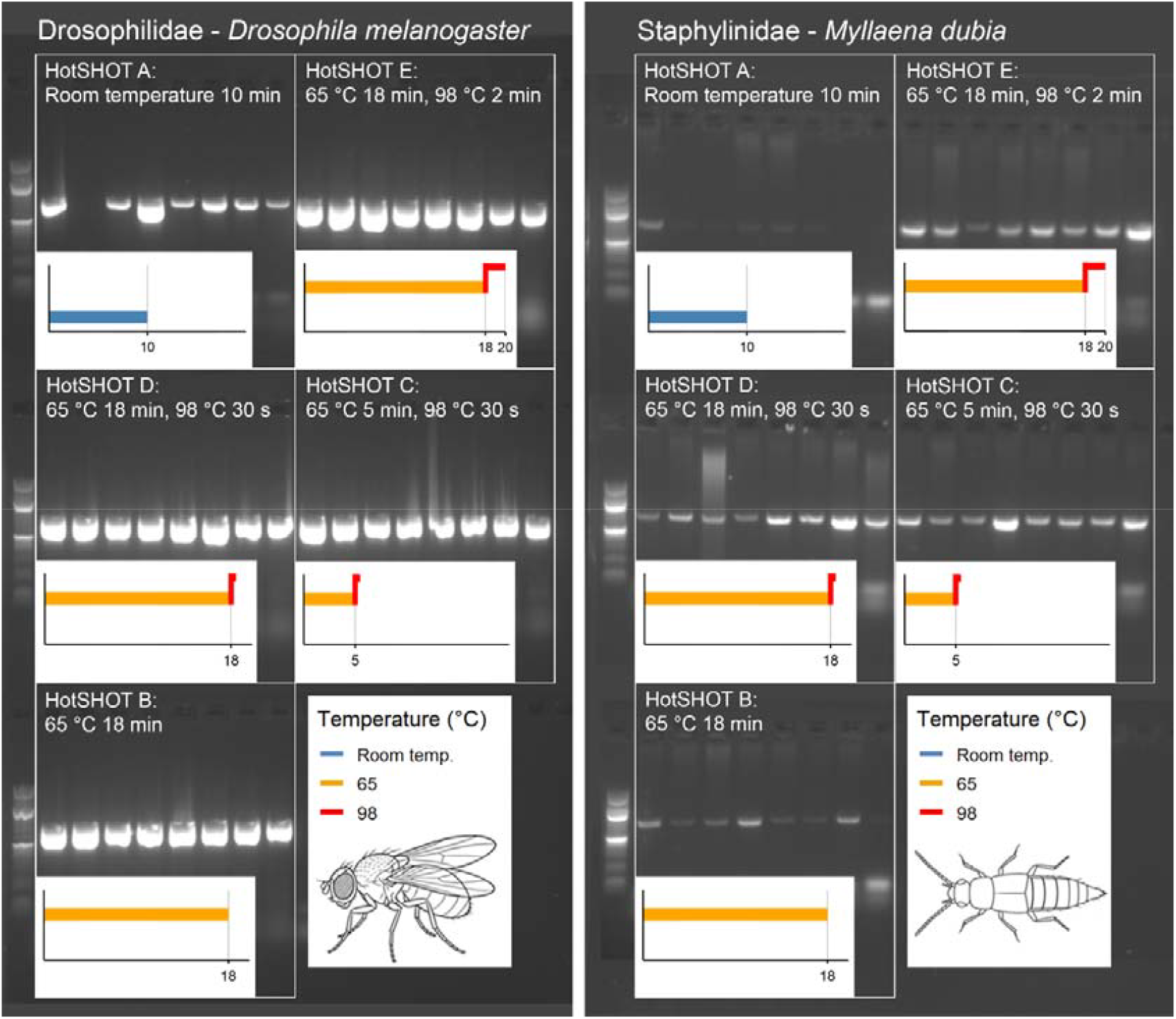
658 bp COI amplification success was high (100% in both taxa except “A”) across all HotSHOT treatments used. Note the fainter gel electrophoresis bands for *M. dubia* specimens treated without any 98 °C step.

## Discussion

Our results show that HotSHOT-treated specimens retain sufficient gDNA for downstream genomic analysis. Voucher gDNA quantities (11.78 – 6,000 ng) are well within the input requirements of commonly used Illumina library preparation kits (e.g. Illumina DNA Prep: 1 – 500 ng input, QIAseq FX DNA Library Kit: 20 pg – 1 µg input, NEBNext Ultra II FS DNA Library Prep Kit: 100 pg – 500 ng input). In some cases, the recovered amounts also fell within the nominal range for long-read sequencing kits such as the Oxford Nanopore Ligation Sequencing Kit V14, but whether DNA exposed to HotSHOT is truly suitable for generating high-quality long-read data will require testing. Taken together, our results suggest that a two-step workflow in which HotSHOT is first used to obtain DNA barcodes for sorting into molecular operational taxonomic units (mOTUs) leaves enough gDNA of sufficient quality for genomics as long as the data are generated with short-read technologies. It thus also appears realistic to use the workflow to address not only the Linnean, Wallacean, and Prestonian but also the Darwinian, Raunkiæran, Hutchinsonian, and Eltonian shortfalls as envisioned by recent reviews of the use of genomics in biodiversity science (Chua et al., 2023; Meier et al, 2025b).

Our data confirms that gDNA integrity in insect specimens is strongly determined by preservation and storage history. The gDNA of Malaise trap samples exposed to tropical field conditions already has heavily fragmented DNA ranging from 300 bp to 5 kbp. Additional HotSHOT treatment thus inflicts limited further damage to DNA in the > 5 kbp size range category. Similar patterns are well documented in museomics, where sequencing success and recoverable fragment length overall decline with specimen age and depend strongly on preservation method, yet low-coverage genomic sequencing has become feasible even for material that is decades to more than a century old (e.g. Strutzenberger et al., 2012; Prosser et al., 2016; Gilbert et al., 2007; Tin et al., 2014). The data suggest that a few weeks of exposure in a Malaise trap in tropical temperatures mirrors the gradual DNA degradation over considerably longer time observed in specimens from natural history museums. However, as in museomics, this level of degradation does not preclude short-read based genomic analysis, but it does severely limit the usability of such samples for long read-based analyses, such as *de novo* genome assembly.

Since the level of DNA degradation limits the sequencing options, two-step workflows that aim to maximize information from bulk biodiversity samples should minimize damage in the first DNA leaching step. Our results show that here HotSHOT treatment choice matters: the harshest but also most popular treatment released the most DNA but at the cost of severely reduced integrity, while milder treatments such as “B” yielded sufficient DNA for barcoding while preserving longer fragments useful for genomic work. Comparative studies of extraction methods also indicate that HotSHOT typically yields very low concentrations and purity compared to column-based protocols (Holmquist et al., 2025; Martincová and Aghová, 2020), which is consistent with our finding that the amount of DNA removed by HotSHOT is so small that it does not preclude obtaining significant amounts of DNA in subsequent genomic extractions (see Fig. 1). Even the most aggressive treatment that maximizes DNA yield in HotSHOT leads only to a modest and not-significant reduction in DNA remaining in the specimen. This bodes well for obtaining genomic data for voucher specimens that had been barcoded.

Our results show a significant interaction between taxon and treatment in experiment 1, with Drosophila yielding more and higher-integrity DNA than Myllaena under mild HotSHOT conditions. These differences cannot be attributed to a single factor, since extraction success is likely determined by multiple specimen properties, including the total amount of DNA available and its accessibility, for which sclerotization may be a proxy. A universal HotSHOT protocol will therefore not be optimal for all specimens. Sorting trap catches by factors such as sclerotization would make sense, but it would also be expensive unless automation is embraced. Robotic sorting systems such as the DiversityScanner, which combines high-throughput handling with convolutional neural network– based family-level classification (Wührl et al., 2022), could provide the technical foundation for automating these choices. This would include the development of predictive AI models that assign specimens to optimal treatments based on traits that explain variation in DNA yield. Such models should also consider the amount of genomic DNA remaining in the specimen to balance barcoding efficiency with preservation for downstream genomic analyses.

In summary, sorting specimens into putative mOTUs based on barcodes directly addresses three major biodiversity shortfalls, namely the Linnean, Wallacean and Prestonian shortfalls, while the remaining Darwinian, Raunkiæran, Hutchinsonian and Eltonian shortfalls can be addressed by genomic sequencing of vouchers. The latter could be further optimized by reducing the damage to DNA incurred in the field and by tailoring HotSHOT treatments to what taxon is being extracted. Shorter exposure of specimens to field conditions and automation provide realistic paths toward such individualized treatments. Despite the modest sample size and single field condition tested, we believe our results provide actionable evidence that HotSHOT is well suited for a two-step approach to studying bulk biodiversity samples holistically.

## Acknowledgements

A.B. and C.H.G. were supported by the Czech Science Foundation grant Junior STAR No. 23-08010M.

## Author Contributions

A.B. and R.M. conceived the study. V.F. and C.H.G. carried out specimen collection and laboratory work. V.F. performed data analyses. V.F. and R.M. wrote the manuscript with input from all authors. All authors read and approved the final version of the manuscript.

## Data Accessibility

All data generated is available from https://doi.org/10.6084/m9.figshare.30265897.v1.

## Benefit-sharing Statement

Benefits from this research accrue from the sharing of our data and results on public databases as described above.

## Notes

https://doi.org/10.6084/m9.figshare.30265897.v1

### Competing Interest Statement

The authors have declared no competing interest.

